# The genome of *Lolium multiflorum* reveals the genetic architecture of paraquat resistance

**DOI:** 10.1101/2024.01.02.573904

**Authors:** Caio A. Brunharo, Aidan W. Short, Lucas K. Bobadilla, Matthew A. Streisfeld

**Author notes:** Author for correspondence: Caio Brunharo, 160 Curtin Rd, University Park, PA, USA.

## Abstract

- Herbicide resistance in agricultural weeds has become one of the greatest challenges for sustainable crop production. The repeated evolution of herbicide resistance provides an excellent opportunity to study the genetic and physiological basis of the resistance phenotype and the evolutionary responses to human-mediated selection pressures. *Lolium multiflorum* is a ubiquitous weed that has evolved herbicide resistance repeatedly around the world in various cropping systems.
- We assembled and annotated a chromosome-scale genome for *L. multiflorum* and elucidated the genetic architecture of paraquat resistance by performing quantitative trait loci analysis, genome-wide association studies, genetic divergence analysis, and transcriptome analyses from paraquat-resistant and -susceptible *L. multiflorum* populations.
- Results suggested that two regions of chromosome 5 were associated with paraquat resistance. The regions contain candidate genes that encode cellular transport functions, including a novel multidrug and toxin extrusion (MATE) protein, and a cation transporter previously shown to interact with polyamines.
- Our results reveal the genetic architecture of paraquat resistance and identified promising candidate genes for future functional studies. Given that *L. multiflorum* is a weed and a cultivated crop species, the genomic resources generated will prove valuable to a wide spectrum of the plant science community.

## INTRODUCTION

Agricultural activities have tremendously modified the landscape and ecological dynamics of its original inhabitants (Tilman, 1999). The need to increase food production over the past 50 years has led to a drastic simplification of agroecosystems (Pretty, 2018). Weed species have colonized these highly disturbed areas and now compete with crops for resources. Chemical control with herbicides is the main strategy used in modern agriculture, and overreliance on these agents has resulted in the widespread evolution of herbicide resistance (Heap, 2023). The evolution of herbicide resistance is one of the greatest challenges to sustainably produce food, fibers, and fuel (Peterson et al., 2018).

Much remains unknown about the physiological and evolutionary mechanisms that lead to the evolution of herbicide resistance. Resistance mechanisms in weeds are classified in two categories: target-site and non-target-site (Gaines et al., 2020). The genetics of target-site resistance has been studied extensively and involves alterations to the herbicide’s target-site. By contrast, the genetic basis of non-target-site resistance is believed to be more complex (Suzukawa et al., 2021). Although it has been shown that non-target site resistance can be caused by increased herbicide metabolism (Brunharo et al., 2019) or sequestration of the herbicide to the vacuole (Brunharo & Hanson, 2017), the genetic controls remain poorly understood. Indeed, basic questions about the genetic architecture of non-target site resistance, including the number of loci affecting the trait, their distribution across the genome, dominance relationships, and whether they involve changes in gene expression, often remain unanswered. Herbicide resistance provides an excellent opportunity to study the evolutionary responses to human-mediated selection pressures.

*Lolium multiflorum* Lam. is a winter annual species native to Europe, temperate Asia, and Northern Africa, but human activities over the past 200 years have caused its spread to all continents (Humphreys et al., 2009). It is a troublesome weed in agriculture, where it can cause drastic reductions in yield if not controlled (Appleby et al., 1976). Management of *L. multiflorum* is primarily performed with herbicides, and the recurrent use of a few herbicide chemistries has resulted in the evolution of >70 herbicide-resistant populations across 14 countries.

Paraquat is an herbicide discovered in 1955 (Brian et al., 1958) that has been used extensively around the world for non-selective weed control. It continues to be one of the most widely used herbicides in the United States (EPA, 2023). Paraquat is actively taken up by plasma membrane-localized transporters, where it must reach its target site in the chloroplasts (Hart et al., 1992a). Once absorbed, it inhibits photosystem I by functioning as a preferential electron acceptor, diverting electrons from ferredoxin to O_2_ and generating reactive oxygen species (ROS) (Hawkes, 2014). In plants susceptible to paraquat, this fast reaction overwhelms the endogenous ROS quenching mechanisms, leading to membrane peroxidation and cell death within hours of treatment (Bromilow, 2004).

Restricted paraquat movement has been reported in resistant weed populations (Brunharo & Hanson, 2017, Soar et al., 2003, Yu et al., 2010), which has been attributed indirectly to enhanced vacuolar sequestration of the herbicide by tonoplast-localized transporters (Brunharo and Hanson, 2017). Enzymes in the Halliwell-Asada cycle are involved in ROS quenching, and their increased activity can provide paraquat resistance. In *Conyza bonariensis*, for example, increased expression of some of these enzymes was observed in resistant individuals (Ye & Gressel, 2000).

Although much remains unknown about the genetic and physiological mechanisms that evolved in natural weed populations, some insight into paraquat resistance mechanisms has been gained by studying *Arabidopsis* mutants (reviewed by Nazish et al., 2022). Given that paraquat’s target site is localized in the chloroplasts, mechanisms that prevent or reduce its movement can limit its action. Xi et al. (2012) identified a loss-of-function mutation in *pqt24-1*, a gene that encodes a plasmalemma-bound ATP binding cassette (ABC), exhibiting reduced cell influx of paraquat. A knockout of *mrv1* (encodes a polyamine transporter) reduced cellular paraquat uptake (Fujita et al., 2012). Restriction of paraquat trafficking from the Golgi apparatus to chloroplasts has been observed in *par1* mutant lines (Li et al., 2013), which contain a non-synonymous mutation in the gene that encodes *AtLAT4*, another member of the polyamine transporter superfamily. Enhanced paraquat tolerance can also be conferred by enhanced vacuolar sequestration and cellular efflux. An amino acid substitution in *DETOXIFICATION EFFLUX CARRIER* (*DTX6*), a member of the multidrug and toxic compound extrusion (MATE) family, was suggested to increase affinity to paraquat (Lv et al., 2021b). It should be noted that the doses of paraquat used in *Arabidopsis* mutants can be six orders of magnitude lower than those that resistant weeds can withstand (0.1 µM in tolerant *Arabidopsis*, 0.13 M in paraquat-resistant *L. multiflorum*; (Brunharo & Hanson, 2017, Fujita et al., 2012), potentially suggesting that different mechanisms could be at play in naturally-evolved weed populations.

Resistance to multiple herbicides in *L. multiflorum* has recently been documented from agricultural fields in the western US (Brunharo & Tranel, 2023, Brunharo & Streisfeld, 2022, Bobadilla et al., 2021, Brunharo & Hanson, 2018), with clear evidence of widespread gene flow among populations, as well as repeated, independent herbicide resistance evolution. However, genetic resources in *L. multiflorum* remain limited. A draft genome from a forage variety of *L. multiflorum* has been created but remains highly fragmented (Copetti et al., 2021). Therefore, a more contiguous, chromosome-level assembly would be an essential resource for dissecting the genetic basis of important traits, such as herbicide resistance. In this context, the three primary objectives of this study are to 1) assemble and annotate the first, full chromosome-level reference genome for *L. multiflorum*, 2) elucidate the genetic architecture and gene expression changes associated with paraquat resistance in *L. multiflorum*, and 3) identify the evolutionary signatures of human-mediated selection pressure in the genome. To do so, we use genetic mapping, a genome-wide association study (GWAS), transcriptome analyses, and population genomic scans for signatures of recent selection between resistant and susceptible populations. Elucidating the genetic architecture responsible for herbicide resistance can provide insights into how organisms respond to strong selection pressure, and it can help initiate efforts to improve weed management practices in the long term.

## MATERIAL AND METHODS

### *Lolium multiflorum* genome assembly and annotation

A *L. multiflorum* individual was grown from seed in autoclaved sand from a previously characterized susceptible population (population Gulf; Brunharo & Streisfeld, 2022) and hydroponically irrigated with half-strength Hoagland’s solution. A tiller was clone-propagated to a larger pot filled with potting soil and grown to maturity. Leaf tissue was collected from plants for high-molecular weight DNA extraction (Wizard HMW DNA Extraction, Promega, Madison, WI). We used flow cytometry to estimate genome size relative to *Conyza canadensis* and *Solanum lycopersicum*. Genomic DNA was sheared with a Megaruptor 3. Sheared DNA was converted to a sequencing library with the SMRTbell Express Template Prep kit 3.0, prior to sequencing on 11 SMRTcell 8M for generation of long-read PacBio HiFi reads on a Sequel IIe platform with 30 h movie time. The combined yield was 244 Gb of raw HiFi data. RNA was extracted from root, leaf, pistil, and anther tissue with the RNeasy Plant Mini Kit (Qiagen, Germantown, MD), and individual libraries were generated with the Iso-Seq SMRTbell prep kit 3.0, generating 23 Gb of HiFi data. To improve the assembly, we used proximity ligation (Hi-C) to capture the 3D structure of chromosomes with the restriction enzymes DpnII, DdeI, HinFI, and MseI and generated 138 Gb of Illumina paired-end sequences.

We generated an initial haploid assembly using *Hifiasm* (v. 0.19.2) (Cheng et al., 2021). HiFi reads were assembled in the integrated mode along with the Hi-C data. This initial assembly was further refined with the *purge_dups* pipeline. Scaffolding was performed with *YaHS* (Zhou et al., 2022) after processing of HiC reads, following the Arima genomics pipeline (https://github.com/ArimaGenomics/mapping_pipeline). Manual curation was performed with *Juicebox* (Durand et al., 2016). Finally, we conducted another round of scaffolding with *RagTag* (Alonge et al., 2022) using the *scaffold* module to order scaffolds onto the genome assembly of the closely related *L. perenne* (Frei et al., 2021).

Full-length transcript sequences were individually processed following the *Iso-Seq* and *tama* pipelines (Kuo et al., 2020). Briefly, sequencing primer removal and demultiplexing was performed with *lima*, poly(A) tail trimming and concatemer identification and removal with *refine*, and clustering of reads and polishing with *cluster.* Processed reads from each tissue were aligned to the *L. multiflorum* reference genome with minimap2 (Li, 2018) and individually collapsed with *tama_collapse*, followed by merging with *tama_merge*.

Repetitive elements were identified with *EDTA* (v2.1.0; Ou et al., 2019). The coding sequences generated in the previous step were provided to *EDTA* to improve repetitive element detection. *RepeatModeler* (Flynn et al., 2020) was also used to identify any remaining transposable elements missed by EDTA.

Genome annotation was performed with *Maker* (Campbell et al., 2014). We included the merged transcripts generated from the *Iso-Seq/tama* pipeline, as well as protein sequences from *Brachypodium distachyon*, *Hordeum vulgare*, and *L. perenne* obtained from Ensembl Plants for annotation based on protein homology. Repeats were masked with RepeatMasker, including the species-specific library created with *EDTA*. We employed Augustus (Stanke et al., 2006) and SNAP (Korf, 2004) *ab initio* gene predictors to train and predict genes, in addition to transcript, protein, and repeat alignments. Functional annotation was performed by querying the protein dataset generated against the Uniprot and interproscan databases (Jones et al., 2014).

We studied the phylogenetic relationships of *L. multiflorum* with other related taxa in the Poaceae family. We used *OrthoFinder* (v2.5.4; Emms & Kelly, 2019) to identify single-copy orthologs from *L. perenne*, *B. distachyon*, *Triticum aestivum*, *H. vulgare*, *Setaria viridis*, *Echinochloa crus-galli*, and *Oryza sativa*, and we produced alignments with *MAFFT* (Katoh et al., 2002). A phylogenetic analysis was performed with *RAxML-NG* (Kozlov et al., 2019), and plotted with *ggtree* (Xu et al., 2022) and treeio (Wang et al., 2019). We used the CoGe (https://genomevolution.org/coge/) platform to obtain the pairwise synonymous mutation rates (K_s_) between *L. multiflorum* and *L. perenne*, *H. vulgare*, *T. aestivum*, and *B. distachyon*. Divergence time was calculated based on a mutation rate of 5.76174 x 10^a−9^ (De La Torre et al., 2017).

### Quantitative trait locus (QTL) mapping of paraquat resistance

We used QTL mapping to identify the genomic locations contributing to paraquat resistance. An outcrossed F_2_ population segregating for resistance was generated. To phenotype paraquat resistance, plants from all generations were treated with 1682 g a.i. ha^−1^ of paraquat. Individuals were scored as dead or alive 14 days after treatment. Prior to paraquat treatment, leaf tissue was sampled from individual plants for DNA extraction. We isolated genomic DNA from 47 susceptible and 48 resistant F_2_ individuals and created individually barcoded nextRAD libraries (Russello et al., 2015) and sequenced on a Novaseq 6000 (2 × 150 bp reads).

Paired-end reads were trimmed with *trimmomatic* (Bolger et al., 2014). Reads were aligned to the reference *L. multiflorum* genome with *bwa*, removed PCR duplicates with *samblaster* (Faust and Hall, 2014), and sorted with *samtools* (Li et al., 2009). Sequencing data from resistant and susceptible individuals were pooled separately, and we then used *freebayes* (Garrison and Marth, 2012) to identify SNPs. We used the QTL-seq method, as implemented in *QTLseqr* (Mansfeld and Grumet, 2018), to identify QTL regions associated with paraquat resistance (Takagi et al., 2013). A positive ΔSNP-index that exceeds the 95% confidence interval suggests that identified alleles are significantly associated with the resistance phenotype.

### Resequencing data generation and analysis

While QTL mapping can provide information on the loci involved in the evolution of resistance, the large linkage blocks present in an F_2_ population can make it difficult to narrow down the genomic location(s) involved in the trait of interest. To complement the QTL analysis and to further explore the genetic architecture of paraquat resistance, we performed a GWAS from 94 individuals collected from six populations (two agricultural fields in Oregon and one in California that are resistant to paraquat, two from fields from Oregon that are susceptible, and a known susceptible cultivated variety from Oregon). At the 3-tiller growth stage, plants were sprayed with lethal doses of paraquat (1682 g a.i. ha^−1^) as described above. Survival was recorded two weeks after treatment. Genomic DNA from 94 individuals was sequenced on a Novaseq6000 in 2×150bp mode to generate 10× coverage.

Paired-end sequences were initially processed with *HTStream* (https://github.com/s4hts/HTStream). Processed files were aligned to the *L. multiflorum* genome with the *minimap2* module for short-read sequences, and PCR duplicates were removed with the *MarkDuplicates* tool of *gatk* (Poplin et al., 2018). Variant detection was performed with *freebayes* (Garrison & Marth, 2012). We used *bcftools* (Danecek et al., 2021) to retain biallelic variants with depths between 10 and 250 in at least 75% of the samples. To obtain an overview of the structural genetic diversity among *L. multiflorum* populations, we identified small variants with *manta* (Chen et al., 2016). Genome-wide association analysis was performed with *GAPIT* (Lipka et al., 2012), with the Enriched Compressed Mixed Linear Model (ECMLM) (Li et al., 2014) and correcting for population structure with PCA eigenvalues and a kinship matrix. Upon identification of statistically significant markers, annotated genes within 2 Mb upstream and downstream of the marker with lowest significance were identified with *bedtools* intersect (Quinlan & Hall, 2010).

### RNA-seq data generation and analysis and weighted correlation network analysis

To assess the effects of differences in transcription level conferring paraquat resistance, we compared gene expression levels among individuals from two independently generated F_3_ populations segregating for paraquat resistance that originated from population PRHC (Brunharo & Hanson, 2018) and population pop60 (Bobadilla et al., 2021). At the 2-leaf growth stage, plants were treated with a lethal rate of paraquat (1682 g a.i. ha^−1^). Leaf tissue was collected from independent F_3_ plants at multiple time points: 0 (immediately before application), 3, 6, 12, and 24 hours after treatment. Tissue was snap frozen in liquid nitrogen. At each timepoint, leaf tissue was collected from four resistant and four susceptible individuals from each of the F_3_ populations. Plants were scored 7 d after application as dead or alive, and chlorophyll fluorescence was measured with a portable fluorometer (OS1p+, Opti-Sciences, Hudson, NH) at each collection time. RNA was extracted from samples with a commercial kit (RNease Plant Mini Kit), and 3’-Tag-RNA-seq sequencing libraries were generated with the QuantSeq FWD kit (Lexogen GmbH, Vienna, Austria). Sequencing was performed on an Illumina Novaseq 6000 in 2 × 150bp. Forward sequences were filtered using *BBDuk* (https://sourceforge.net/projects/bbmap/). Filtered reads were aligned to the reference genome with *STAR* in *quantMode* (Dobin et al., 2012). Differential gene expression was quantified with the R package *edgeR* (Robinson et al., 2009) and *limma* (Ritchie et al., 2015).

To determine if differentially expressed genes between resistant and susceptible individuals tended to be co-expressed with genes with similar functions, we performed a Weighted Co-Expression Network Analysis (WGCNA) with the *WGCNA* package (Langfelder & Horvath, 2008) using the chlorophyll fluorescence as the response variable. To identify any outlier samples in our dataset, we employed hierarchical clustering, and subsequently used the constant-height tree-cut function to eliminate the outliers. The appropriate soft-threshold power was identified by performing the approximate scale-free topology criterion. We then derived a signed adjacency matrix through bi-weight mid-correlation and a signed topological overlap matrix through dissimilarity calculations. Genes were grouped into modules using hierarchical clustering, and we employed the dynamic tree-cutting algorithm to partition genes into distinct modules. Next, we computed module eigengenes, which allowed us to merge similar modules and pinpoint modules associated with paraquat resistance. To identify genes that exhibit a strong correlation with genes in modules linked to paraquat resistance (i.e., hub genes), we conducted an intra-modular analysis. Hub genes were identified based on their module membership (ranging from 0 to 1, indicating overall connectivity) and their gene-trait significance (determined by the Pearson correlation between expression and the trait). Finally, to gain insights into the biological functions associated with these significant modules, we conducted a Gene Ontology (GO) term enrichment analysis for each of them using *TopGO* R package.

### Population genomic analyses of selection on resistance

To complement the genetic mapping approaches described above, we used population genomic information from the different susceptible and resistant populations to determine the role of human-mediated selection on the evolution of paraquat resistance. The application of paraquat in agricultural fields is expected to confer a strong selective pressure for resistance, resulting in signatures of a selective sweep surrounding the loci involved in resistance. However, if the genetic variation responsible for resistance existed in the population prior to the onset of paraquat application, functional variants would likely occur on multiple haplotypes, which might obscure the signatures of selection. We calculated levels of genetic divergence (Fst) between the resistant and susceptible populations across the genome, with the expectation that locally elevated patterns of genetic divergence between susceptible and resistant populations would be associated with resistance. We estimated Fst using the Weir method (Weir and Cockerham, 1984) at each biallelic SNP after removing indels, as implemented in *VCFtools* (Danecek et al., 2011). These were plotted across the genome, and regions of exceptionally elevated Fst were considered as potentially selected loci.

We also asked if the three resistant populations we sampled were differentiated in the same regions of the genome. Elevated Fst could be caused by only one or two resistant populations being differentiated from the susceptible populations, but this would not indicate a region that is always involved in resistance. To test this, we calculated Fst in each 25 kb window across the genome between each pair of resistant and susceptible populations using the *popgenwindows.py* script (https://github.com/simonhmartin/genomics_general). For each of the nine pairwise comparisons, we counted the number of times each window was found in the top 1% of windows across the distribution of Fst values, because the more often a window is found in the top 1% of Fst values across comparisons, the more likely the same resistance haplotype is diverged from susceptible haplotypes.

Fst can be considered as a relative measure of sequence divergence because its value is influenced by levels of genetic diversity within populations (Cruickshank & Hahn, 2014). Although a region of locally elevated Fst could be caused by reduced diversity in either (or both) susceptible or resistant groups, we expect that selection occurred in resistant individuals. Therefore, we expect to find locally reduced genetic diversity and fewer segregating sites (i.e., polymorphism) in resistant relative to susceptible individuals. In addition, we expect resistant individuals to show a higher average frequency of alternate alleles relative to the reference. The reference assembly is derived from a susceptible individual, so a higher frequency of alternate alleles in resistant individuals would be consistent with selection increasing the frequency of alleles responsible for resistance. Based on overlap between the genetic mapping and Fst results, we focused these analyses on two regions on chr5 (see Results). Finally, we calculated the site-specific extended haplotype homozygosity statistic. Due to a recent selective sweep, a rapid increase in the frequency of a beneficial mutation will result in elevated linkage disequilibrium, leading to extended patterns of homozygosity within haplotypes (Sabeti et al., 2002, Smith & Haigh, 1974, Voight et al., 2006). However, in the absence of selection, we expect haplotypes to break down over time due to new mutations and recombination. We compared haplotypes from susceptible and resistant individuals by focusing on two sites that had the highest Fst in each of the two regions on chr5. Haplotype and marker information was extracted from each VCF file using the *data2haplohh* function of the *rehh* package (Gautier & Vitalis, 2012). EHH was calculated from the marker and haplotype information for each of the taxa using the *calc_ehh* function in *rehh*.

## RESULTS

### A chromosome-level assembly and annotation for *L. multiflorum*

The first step towards elucidating the evolution of herbicide resistance in *L. multiflorum* was to assemble a high-quality reference genome for this species (Fig. 1). We generated 45× coverage long-read data from DNA of an herbicide susceptible individual and used proximity ligation (Hi-C) to generate a haploid assembly of the *L. multiflorum* genome (Table S1). The *de novo* assembly of the *L. multiflorum* genome resulted in 297 scaffolds, with N_50_ = 61 Mb and L_50_ = 12 (i.e., the length sum of 12 contigs contributes to at least 50% of the assembly). This initial assembly was improved by ordering scaffolds based on the synteny shared with the closely related species *L. perenne*, resulting in 7 chromosomes containing 2.55 Gb of nuclear content (90% of the genome; N_50_ = 363 Mb; Table S1). The genome size obtained *in silico* supports the flow cytometry estimates of 2.72 Gb haploid size. BUSCO analysis indicated that 93% of the single-copy orthologs from the *poales_odb10* dataset were contained in the assembly. Approximately 83% of the DNA content is composed of repetitive elements (Table S2), as expected from a plant species with a large genome size (Schnable et al., 2009). We annotated the *L. multiflorum* genome with 23 Gb of Iso-Seq data from leaf, root, pistil and anther tissue and identified 49,295 protein-coding sequences.

**Fig. 1.**
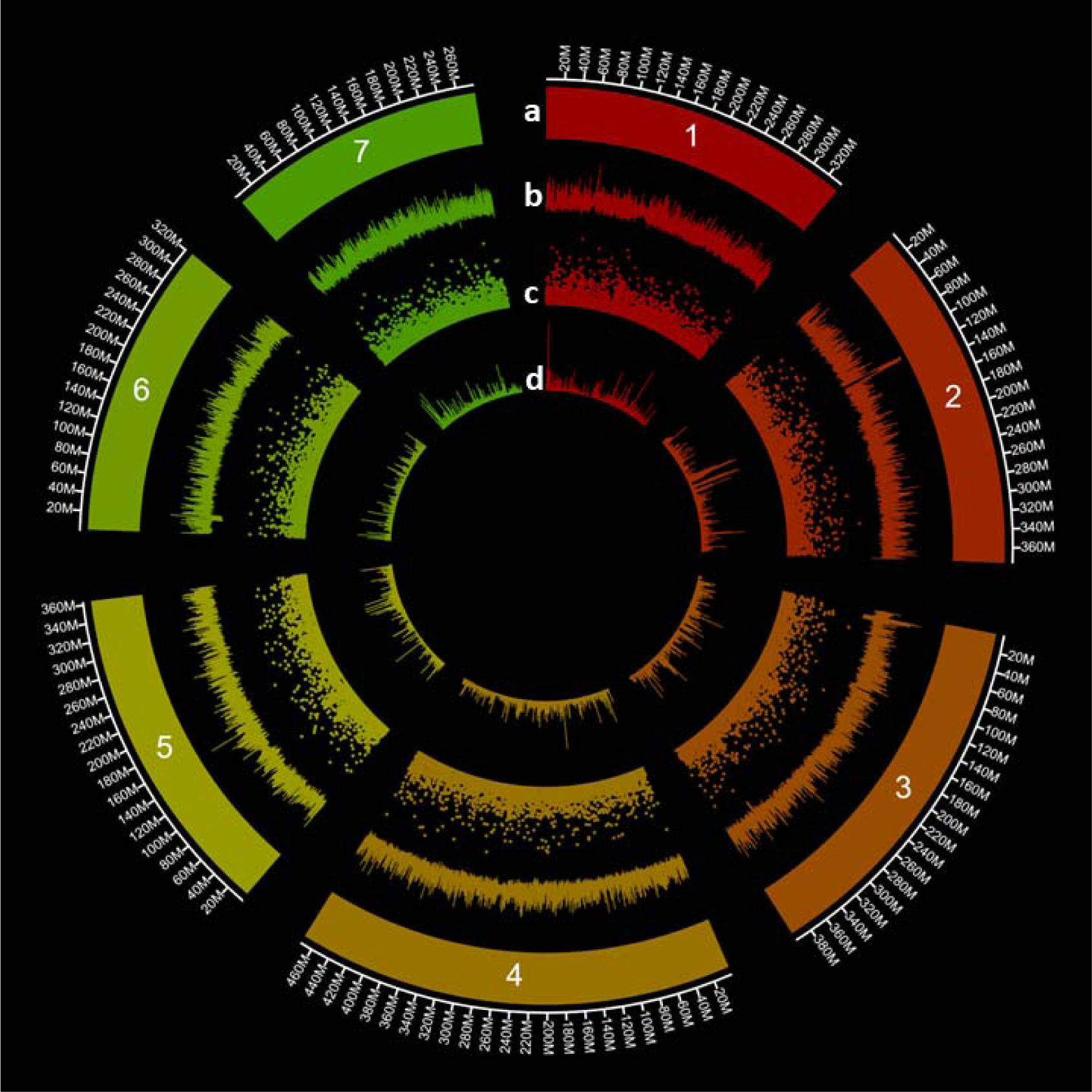
Circos plot of the *L. multiflorum* genome and its various features, plotted in 100 kb windows, where peak height represents feature frequency. Tracks represent a) the seven chromosomes, b) repetitive elements, c) small variants (<1000 bp), and d) single-nucleotide polymorphisms.

In weed management, accurate species identification is crucial, because different species can exhibit distinct responses to efforts to control them. *Lolium multiflorum, L. perenne*, and *L. rigidum* are often referred to, or treated as, a single species for basic biology and management (Scarabel et al., 2020, Yanniccari et al., 2020). Our results confirm these species are closely related, but they are phylogenetically distinct, confirming morphological and life-history studies performed elsewhere (Fig. S1A) (Bararpour et al., 2017). In addition, based on the synonymous substitution rates (K_s_) between homologous gene pairs, we found that *L. multiflorum* appears to have diverged from *L. perenne*, *T. aestivum*, *H. vulgare*, and *B. distachyon* approximately 5.4, 26, 27, and 29.4 M years ago (Fig. S1B).

### Genetic mapping reveals the genetic architecture of resistance

A QTL analysis in an F_2_ mapping population identified regions of the genome segregating with the resistance trait (Fig. 2). We identified two regions on chr5 that were significantly associated with paraquat resistance based on the ΔSNP-index exceeding the 95% confidence interval. The intervals of the two detected QTL spanned positions 120,860,887 - 167,802,303, and 294,188,538 - 363,361,071 on chr5. This could suggest two separate loci on chr5 are responsible for paraquat resistance, but further analysis is necessary to determine the precise regions involved.

**Fig. 2.**
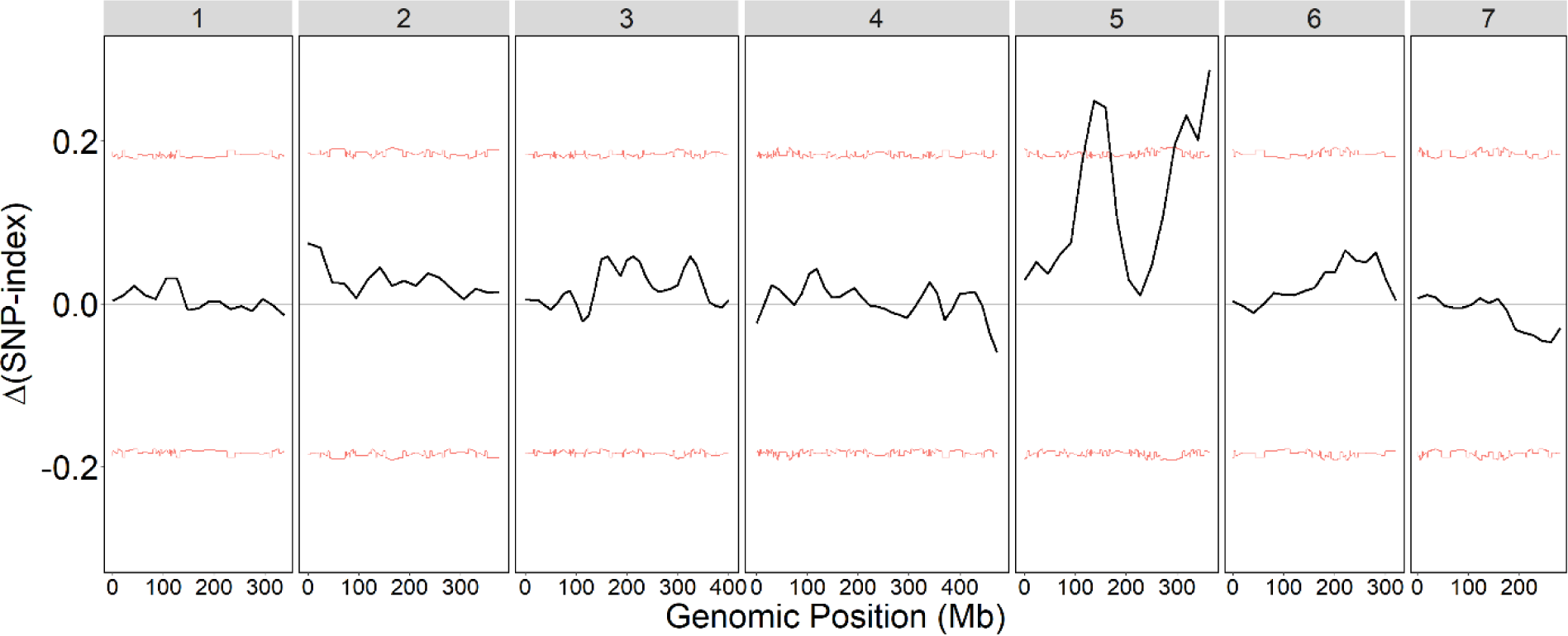
QTL-seq analysis from an F_2_ mapping population between paraquat-resistant and - susceptible *L. multiflorum* individuals. The numbers at the top represent the chromosome number. The red lines represent the 95% bootstrap confidence intervals based on 10,000 replicates used for significance testing.

To further explore the genetic architecture of resistance, we performed a GWAS by resequencing 94 *L. multiflorum* individuals to an average 10× coverage, and implemented statistical models in *GAPIT* (Lipka et al., 2012) to identify SNPs and INDELs associated with paraquat resistance. GWAS can take advantage of the genetic variation present across multiple individuals that have a longer history of recombination. Moreover, determining the genomic regions where the QTL and GWAS analyses intersect would provide independent corroboration of potential candidates for resistance. The final GWAS dataset contained 3,385,096 SNPs and 1,359,667 INDELS, which results in an overall marker density of 1.7 variants per 1 kb. In total, we found 22 markers that were significantly associated with the difference between resistant and susceptible plants (Fig. 3). Although five of these markers were statistically significant, they occurred as singletons, with no additional significant markers in those regions. Therefore, we focused only on the remaining 17 markers that localized to three distinct regions of the genome: one on chr2, and two on chr5. The three significant markers found on chr2 spanned a distance of only 289 bp (between positions 40,259,279 – 40,259,568). Of the remaining significant markers, eight of them localized to a 239 kb region on chr5 (between positions 100,183,991 – 100,423,860). The final five markers that were significantly associated with the phenotype also were located on chr5, between positions 159,409,415 – 165,297,925, a distance of 5.89 Mb. Although the first region on chr5 does not directly overlap with the QTL peaks (roughly 20 Mb away), this second region was contained within the first peak on chr5.

**Fig. 3.**
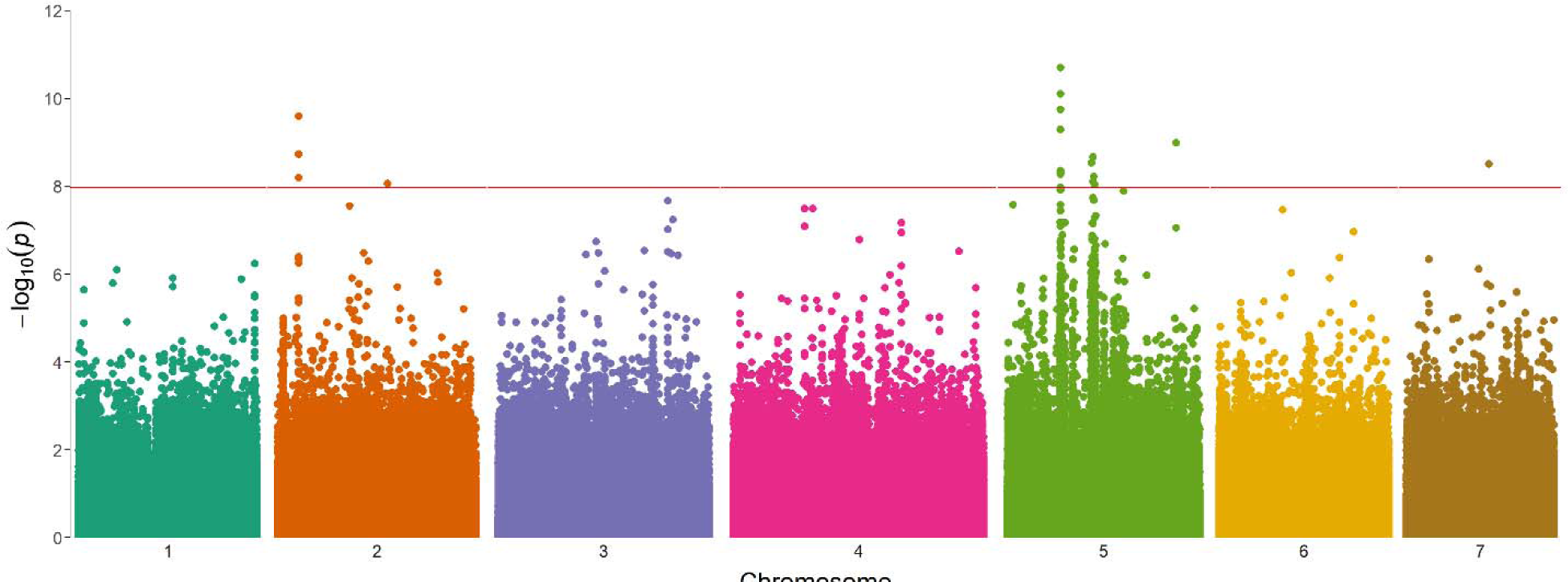
Genome-wide association of paraquat-resistance in *L. multiflorum*. A total of 4,744,764 (n) makers were tested. The red line is the Bonferroni threshold calculated as 0.05/n.

We further investigated these regions by extracting annotated genes in 2 Mb regions surrounding the marker in each region with the lowest P-value (4 Mb total per region). The focal markers were position 40,259,568 of chr2, and 100,269,722 and 162,257,267 of chr5. A total of 214 genes associated with paraquat resistance were annotated near these three regions, of which 89 were on chr2, 64 near the first peak on chr5, and 61 near the second peak of chr5 (Table S3). There were several genes with known functions that could be involved with resistance to paraquat. For example, there were genes that had functions related to the response to oxidative stress, those that mediated intracellular transport, and genes known to respond to herbicides. Genes with potential roles in herbicide metabolism were identified, such as cytochrome P450s, as well as a polyamine oxidase gene that regulates polyamine intracellular concentration. Given that paraquat resistance has been previously suggested to be conferred by reduced herbicide movement, genes that encode membrane-bound transporters are of particular interest. Specifically, we identified *SEC31B* (promotes the formation of transport vesicles from the endoplasmic reticulum) on chr2, *Transmembrane 9 superfamily member 1* (*TMN1*; involved in cell adhesion and phagocytosis in the secretory pathway), *Lysine histidine transporter-like 8* (*AATL1*; a transmembrane amino acid transporter) and *VATP-P1* (a V-type proton ATPase) in the first region of chr5, and *NPF5.10* (*NRT1/PTR FAMILY*, a nitrate or di/tri-peptide transporters), and *DTX10* (a member of the MATE family of proteins) in the second region of chr5.

### RNAseq identifies genes differentially expressed between susceptible and resistant plants

To complement the results from QTL-seq and GWAS, we performed a differential gene expression experiment in two separate F_3_ populations segregating for paraquat resistance. Based on visualization of a multidimensional scaling plot, there was a clear separation between mapping populations. Therefore, analyses were performed separately for each segregating population. A total of 55 genes was differentially expressed in the F_3_ population derived from PRHC, and 10 genes from pop60 (Table S4). Of these, only two genes were found to be differentially expressed in both populations. One of these encodes a chloroplastic uncharacterized aarF domain-containing protein kinase, and the other encodes a glutathione synthetase (GSH2). In both cases, the genes were expressed at higher levels in resistant individuals relative to susceptible, with *GSH2* being the most highly expressed gene in the dataset (Table S4). Interestingly, we found a strong over-representation of differentially expressed genes across chr5. Of the 63 differentially expressed genes detected, 32 (51%) occurred on chr5. Among these, there is only one gene that is differentially expressed and found among the annotated genes near the GWAS hits. This gene encodes a protein with sequence similarity to the NPF5.10 protein from *Arabidopsis thaliana* and is expressed at a higher level in resistant plants from the PRHC-derived population.

### Gene co-expression networks dissect the coordinated expression patterns in response to paraquat

The network analysis grouped genes with similar expression patterns into separate modules, resulting in 18 modules identified in the population derived from pop60 and 28 from PRHC (Fig. S2-3). The number of genes contained within the modules ranged from 74 to 3,288 for pop60, and from 38 to 3,286 for PRHC. Similar patterns were observed across the modules that were positively correlated with chlorophyll fluorescence (a proxy for paraquat resistance). Specifically, these modules contained co-expressed genes with functions that included transmembrane transport, photosynthesis, glutathione biosynthesis, ABC transporters, and superoxide metabolism (Fig. S2-3). Hub genes included detoxification 21 (*DTX21*) and ABC transporters (*ABCB11*, *ABCA8*). In addition, *psbB Photosystem II CP47* and *atpI* ATP synthase subunit α were detected as hub genes in modules associated with photosystem responses and regulation (Fig. S2, module “yellow” and “tan” in pop60 and PRHC, respectively). These results further indicate that membrane-bound transporters are responsive to paraquat application and could be involved in herbicide resistance or to minimize oxidative stress from paraquat application. Of the 60 major hub genes positively correlated with paraquat resistance, 19 (32%) were on chr5. Three annotated genes were found within the QTL interval on chr 5: LOLMU_00024321 (Protein of unknown function), LOLMU_00024251 (Probable protein phosphatase 2C 20), and LOLMU_00027468 (Protein of unknown function).

### Recent positive selection on resistant haplotypes

By scanning the genome for SNPs with elevated Fst, we detected two regions that were highly differentiated between resistant and susceptible individuals (Fig. S4). Importantly, both of these regions occurred on chr5, and they aligned with the two regions on that chromosome identified from the GWAS (Fig. 4). Hereafter, we define these two regions as region 1 and region 2, based on their position on chr5 (region 1: 99.7 Mb, and region 2: 160.1 Mb). Although neither region contained differentially fixed sites (i.e., Fst = 1), Fst did exceed 0.7 at three sites in region 2, indicating extensive sequence divergence between resistant and susceptible populations.

**Fig. 4.**
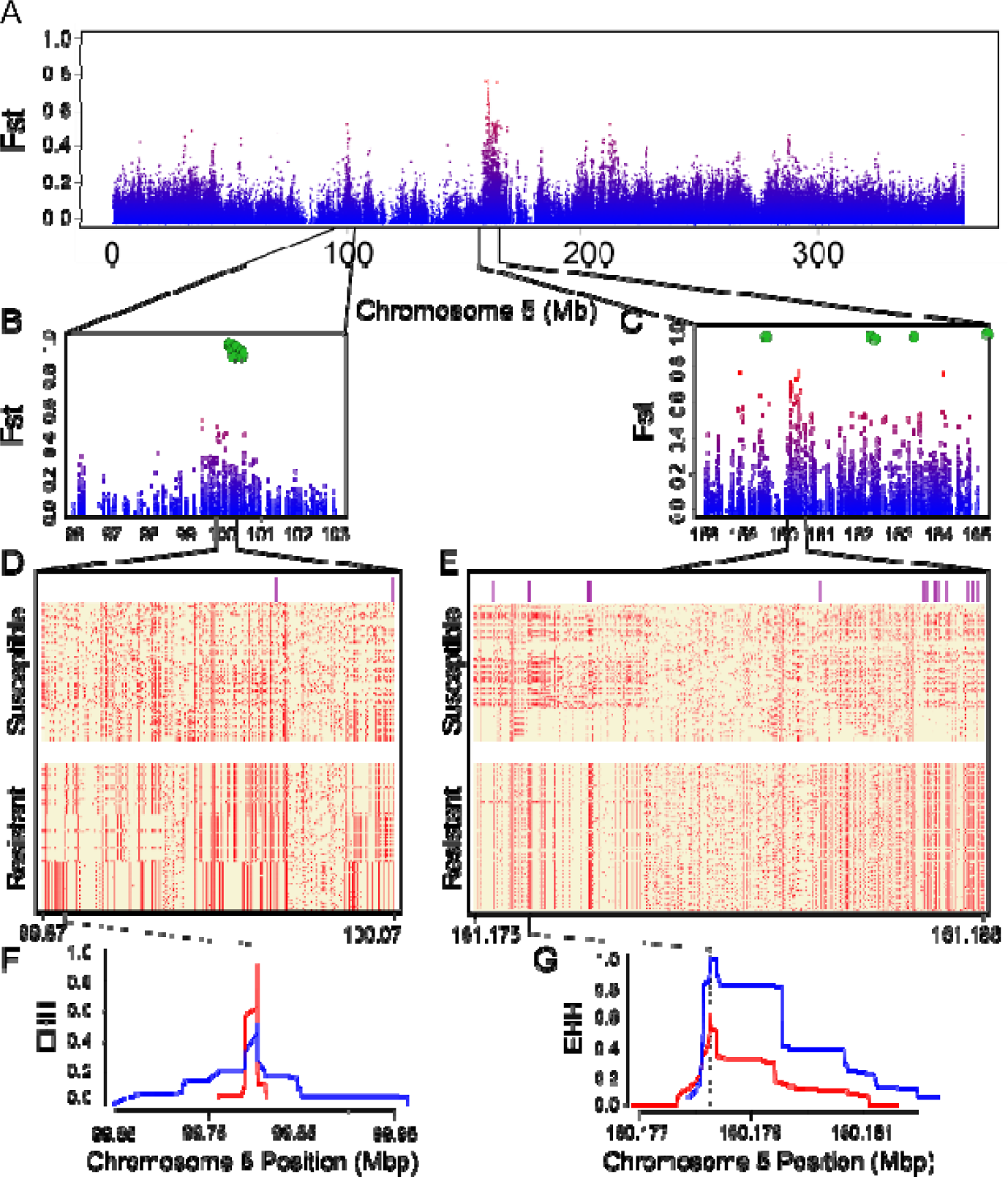
Recent positive selection on resistant haplotypes. A) Fst scan of biallelic SNPs across chromosome 5. The SNPs are color-coded based on their relative values across the entire genome (see Fig. S4), with red indicating higher overall divergence. Panels B and C contain the same information as in panel A, but they are zoomed in on regions 1 and 2. Green hexagons indicate the location of GWAS markers that were significantly associated with the resistance phenotype. D, E) Each row is a haplotype from resistant or susceptible individuals, either for region 1 (D) or region 2 (E), and columns correspond to variant sites. Yellow boxes indicate the presence of the reference allele, while red boxes indicate the alternate allele. Purple lines above the haplotypes denote the sites where Fst between resistant and susceptible populations is greater than 0.4. The range of sites included in each region is reported at the bottom of each panel, in Mb. F, G) The site-specific extended haplotype homozygosity (EHH) statistic for region 1 (F) and region 2 (G), with blue lines corresponding to resistant haplotypes and red lines denoting susceptible haplotypes. The black dotted line indicates the focal SNP included in each analysis (region 1: 99,802,212; region 2: 160,178,236).

Further inspection of these two regions revealed that the divergence between susceptible and resistant populations was caused primarily by a reduction in diversity and polymorphism in resistant, but not susceptible populations (Fig. 4D-E). Haplotype diversity was 45% lower in resistant individuals than susceptible in region 2 (resistant: 5.4 x 10^−4^ ± 2.7 x 10^−5^ S.E.; susceptible: 7.9 x 10^−4^ ± 2.9 x 10^−5^ S.E.). Although diversity was similar between susceptible and resistant populations in region 1 (resistant: 2.8 × 10^−4^ ± 1.9 x 10^−5-^ S.E.; susceptible: 2.9 x 10^−5^ ± 1.6 x 10^−5^ S.E.), the proportion of segregating sites among resistant haplotypes was nearly half that in susceptible individuals in both regions (region 1: resistant = 0.54, susceptible = 0.93; region 2: resistant = 0.61, susceptible = 0.93). These patterns were in stark contrast to the proportion of segregating sites across all chr5, where polymorphism was much more similar between resistant and susceptible populations (resistant = 0.77, susceptible = 0.85). Finally, resistant individuals showed a frequency of the alternate (i.e., non-reference) allele that was nearly twice as high as susceptible individuals in both regions (average frequency among sites for region 1: resistant = 0.22, susceptible = 0.11; region 2: resistant = 0.19, susceptible = 0.10). This difference is even more pronounced when we consider the number of variants where the alternate allele has a frequency greater than 0.5. In region 1, there were 37 sites where the alternate allele was at a frequency > 0.5 in resistant individuals, but only 19 sites in susceptible individuals. In region 2, there were 100 sites in resistant individuals but only 8 among susceptible. The presence of a high frequency of the alternate allele at multiple sites in these regions is consistent with selection increasing the frequency of distinct haplotypes in resistant but not susceptible individuals.

We also detected patterns of extended haplotype homozygosity in both regions that were more pronounced in resistant individuals. In region 1, we found a very broad region of homozygosity (roughly 300 kb) in some, but not all, resistant individuals. This is in stark contrast to the pattern in susceptible individuals, where the signal of EHH is quite narrow (Fig. 4). In region 2, the probability of homozygosity was substantially elevated in resistant individuals relative to susceptible, but this region extended only over 5 kb. Thus, even though the patterns are different in each region, they are both consistent with selection driving an increase in the frequency of the alternate allele.

Finally, we also asked if the same regions of the genome were differentiated in all three resistant populations relative to the susceptible populations. To do this, we calculated Fst in non-overlapping 25 kb windows for all pairs of resistant and susceptible populations and counted in how many of these pairwise comparisons was each window found in the top 1% of the Fst distribution. We found only two regions in the genome where all nine comparisons between resistant and susceptible populations showed the same window in the top 1% of the distribution, and these corresponded to regions within or directly adjacent to regions 1 and 2 (Fig. S5). Indeed, there was a pileup of windows in these two regions where multiple comparisons showed elevated Fst. No other window across the genome exceeded six comparisons where that window was repeatedly in the top 1% of Fst windows.

## DISCUSSION

Herbicide resistance is one of the greatest challenges in sustainable crop production, and elucidating the genetic basis of resistance could aid in improving weed management practices. *Lolium multiflorum* is one of the most troublesome weed species in the world, having evolved herbicide resistance in multiple cropping systems and countries. The genomic resources that we generated in this work will be a valuable tool to dissect the genetic bases of herbicide resistance and other traits. Obtaining a contiguous genome is a crucial component of the assembly effort (Lee et al., 2016) that allows researchers to better understand the genomic context in which genes or variants of interest occur. The *L. multiflorum* genome assembled here had 90% of the DNA content placed on 7 chromosomes (N_50_=363 Mb), with 93% completeness, which is comparable to other high-quality assemblies (Cai et al., 2021, Benson et al., 2023).

The genetic basis of paraquat resistance was addressed using multiple “-omics” datasets. Implementing multiple approaches has proven necessary to dissect the genetic architecture of plant traits (Du et al., 2018, Jiang et al., 2019), as any one experiment may have shortcomings (Li & Xu, 2022). Overall, our QTL-seq, GWAS, and Fst approaches converged on two regions on chr5 that appeared to be associated with paraquat resistance in *L. multiflorum*. GWAS also identified significant SNPs on chr2 and chr7. Although we used a highly conservative Bonferroni test to correct for multiple testing, these associations may represent false positives due to residual population structure, epistasis, or other indirect effects (Platt et al., 2010, Kaler and Purcell, 2019). Furthermore, we found only two regions in the genome where all nine pairwise Fst comparisons between resistant and susceptible populations showed the same window in the top 1% of the distribution, and these corresponded to regions within or directly adjacent to regions 1 and 2 (Fig. S5).

Paraquat resistance in other species, including in the weed *L. rigidum* that is closely related to *L. multiflorum* (Yu et al., 2009), is believed to be caused by a single gene with incomplete dominance. By contrast, (Shaaltiel et al., 1988) suggested the possibility of linkage between two genes involved in paraquat resistance. We found two linked loci that appear to be associated with resistance. One explanation for this finding is that there are genes in regions 1 and 2 that work in concert to generate the resistance phenotype. The resistant populations used in this study were collected from fields that have been subjected to recurrent paraquat applications, resulting in a constant selective pressure. However, recombination between these two regions would break up co-adapted genotypes at these loci. Future experiments with larger sample sizes should explore patterns of linkage disequilibrium between these regions to investigate the possibility of epistatic selection maintaining particular genotypic combinations at these loci.

Analysis of gene expression patterns produced valuable insights into how plants respond to oxidative stress caused by paraquat. When all time points were analyzed together, we observed an overall higher expression of *GSH2* in resistant plants from both F_3_ populations. Closer inspection at each time point provided more information into the temporal dynamics of its expression (Table S4). Notably, we observed that *GSH2* was constitutively expressed at higher levels in the PRHC population (5-fold) prior to herbicide treatment, but this same pattern was not found in the pop60 F_3_ population. However, upon paraquat treatment, *GSH2* expression in resistant individuals increased to 17-fold greater than the susceptible individuals in the pop60 derived F_3_ population. These results are consistent with the different evolutionary histories of pop60 and PRHC (Brunharo and Streisfeld, 2022), which implies that they may have evolved different mechanisms to cope with abiotic stresses. Alternatively, the recent exposure of field populations to paraquat or other herbicides that cause oxidative stress could have induced the constitutive over-expression of oxidative stress-associated genes. By contrast, sublethal oxidative stresses could alter overall gene expression by inducing epigenetic modifications that are trans-generationally stable (Dyer, 2018, Sharma et al., 2022). The WGCNA also found modules with enriched GO terms for hub genes involved in oxidative stress and xenobiotic response that responded to paraquat application.

The population genomic analyses revealed molecular signatures that were consistent with the action of recent positive selection in resistant individuals. Fst analysis identified two regions in chr5 that were highly differentiated between resistant and susceptible populations due to a lower diversity and segregating sites in resistant individuals. We also observed a pileup of windows in these regions where multiple comparisons between resistant and susceptible populations showed elevated Fst (Fig. S5). In addition, a greater number of non-reference polymorphisms was found in the resistant individuals compared to susceptible, which resulted in patterns of extended haplotype homozygosity in resistant plants. When combined with the GWAS and QTL analyses, these genomic data suggest that these regions experienced recent positive selection leading to evolution of resistance.

In addition to revealing the genetic architecture and evolutionary responses to resistance, our extensive genomic analyses reveal promising candidate genes. Specifically, genes that encode NPF5.10 and DTX10 are of particular interest, as both are contained within the GWAS peaks and are adjacent to region 2 from the Fst analysis. NPF5.10 is also the only gene in these regions that was differentially expressed between resistant and susceptible individuals. It is a member of the NRT1/PTR family, which are nitrate and oligopeptide transporters localized to cellular membranes. It has been suggested that an amino acid substitution in the NPF6.4, a member of the NRT1/PTR family, reduced norspermidine (a polyamine) cell uptake in Arabidopsis (Tong et al., 2016). Polyamines are natural compounds responsible for protein synthesis, ion channel modulation and a number of biological processes (Igarashi & Kashiwagi, 2010), and paraquat is known to competitively inhibit polyamine transporters (Hart et al., 1992a, Hart et al., 1992b). (Brunharo & Hanson, 2017) observed that paraquat-resistant *L. multiflorum* pretreated with putrescine (a polyamine) became susceptible upon paraquat application, which suggested that a membrane-bound transporter could be involved in the resistance mechanism.

DTX10 is a member of the MATE family (Multidrug And Toxic compound Extrusion) that is found in prokaryotes and eukaryotes. These transporters are localized to the tonoplast (Zhang et al., 2017), Golgi apparatus (Seo et al., 2012), and plasma membrane (Rogers and Guerinot, 2002). A total of 58 MATE proteins have been annotated in *Arabidopsis* (Hvorup et al., 2003) and 53 in rice (Tiwari et al., 2014), and they have been shown to transport cationic compounds across cellular and organellar membranes (Kusakizako et al., 2020). Most importantly, an amino acid substitution in the DTX6 has previously been shown to confer paraquat resistance in an experimentally-derived *Arabidopsis* accession (Lv et al., 2021a). DTX6 is localized in the endomembrane trafficking system and is responsible for increased vacuolar sequestration and cellular efflux of paraquat (Lv et al., 2021a).

In this work, we elucidated the genetic architecture of paraquat resistance in *L. multiflorum*, and we identified promising candidate genes for future functional validation studies. Understanding the genetic basis of herbicide resistance is crucial to improve the sustainability of cropping systems. A key component of weed management is to limit dispersal of herbicide resistant individuals at multiple levels of the production operation, such as in the field, warehouses, or shipping containers during international trade. This could be accomplished with the development quick resistance identification assays using genetic markers (Brusa et al., 2021), and the development of lateral flow assays that target the genetic variants (Baerwald et al., 2020). The genomic resources generated in this work will be valuable for a wide range of plant scientists. *Lolium multiflorum* is not only an agricultural weed, but domesticated varieties are cultivated as a cover crop and for forage, and the assembled genome could be used by breeders for trait improvement.

## ACKNOWLEDGEMENTS

Funding for this project was provided by the College of Agricultural Sciences at The Pennsylvania State University and Oregon State University.

## AUTHOR CONTRIBUTION

CAB conceived and designed the study, collected data, assembled and annotated the genome, performed QTL, GWAS, RNA-seq, and drafted and revised the manuscript. AWS performed Fst scans and haplotype networks, and revised the manuscript. LKB performed the WGCNA, and revised the manuscript. MAS performed Fst and haplotype networks, and drafted and revised the manuscript.

## SUPPORTING INFORMATION

Fig. S1. Evolutionary relationships between *L. multiflorum* and other species in the Poaceae family. A) Phylogenetic tree of *L. multiflorum, L. perenne*, *Brachypodium distachyon*, *Triticum aestivum*, *Hordeum vulgare*, *Setaria viridis*, *Echinochloa crus-galli*, and *Oryza sativa*. B) Distribution of the synonymous substitution rate (K_s_) between *L. multiflorum* and closely related species.

Fig. S2. Weighted gene co-expression network analysis of the F_3_ paraquat-resistant pop60 population. (A) Heat map of module-trait relationships shows a correlation from more negative (blue) to more positive (red) for each module, which are given names with different colors. Each column indicates a comparison between time points or individuals. (B) Hierarchical cluster trees show the co-expression modules identified by WGCNA.

Fig. S3. Weighted gene co-expression network analysis of the F_3_ paraquat-resistant PRHC population. (A) Heat map of module-trait relationships shows relationship from more negative (blue) to more positive (red) of each module color. Each column indicates a comparison between time points or individuals. (B) Hierarchical cluster trees show the co-expression modules identified by WGCNA.

Fig. S4. Genome scan of Fst at all bi-allelic SNPs between paraquat-resistant and -susceptible populations of *L. multiflorum*.

Fig. S5. The number of pairwise comparisons between resistant and susceptible populations where Fst (calculated in 25 kb windows) was found in the top 1% of the distribution across windows.

Table S1. Summary of *L. multiflorum* assembly and annotation.

Table S2. Repetitive element content in the *L. multiflorum* genome.

Table S3. Genomic regions identified in GWAS and number of annotated genes.

## DATA AVAILABILITY

Raw sequencing data is available at the NCBI Sequence Read Archive under BioProject PRJNA1046158. The genome and annotation files will be available at the Weed Genomics Consortium at https://weedpedia.weedgenomics.org/ upon publication.

