## Supplemental file 1 for "The genome of *Lolium multiflorum* reveals the genetic architecture of paraquat resistance"

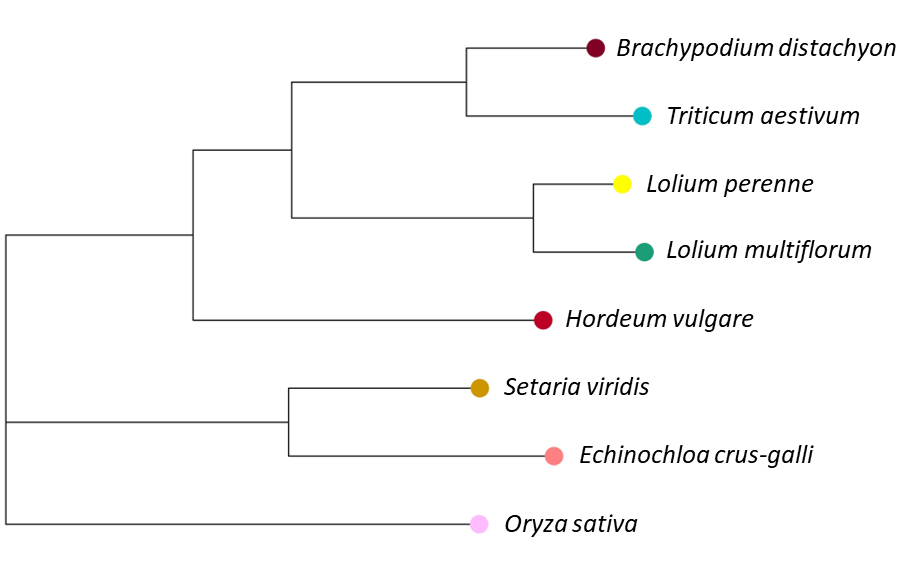

A

B

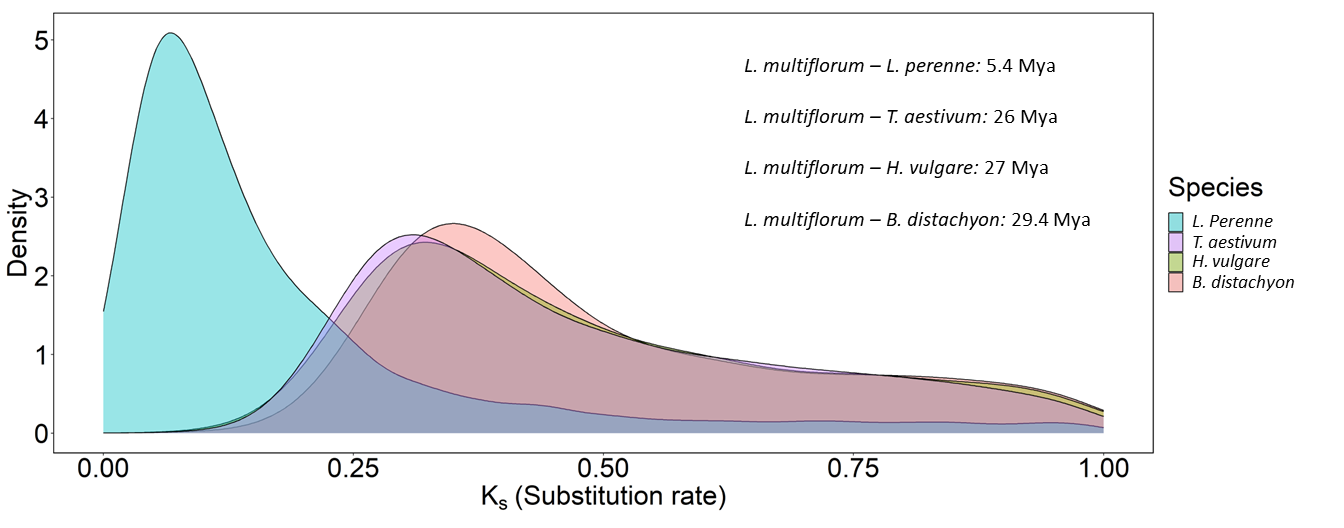

Fig. S1. Evolutionary relationships between *L. multiflorum* and other species in the Poaceae family. A) Phylogenetic tree of *L. multiflorum,* *L. perenne*, *Brachypodium distachyon*, *Triticum aestivum*, *Hordeum vulgare*, *Setaria viridis*, *Echinochloa crus-galli*, and *Oryza sativa*. B) Distribution of the synonymous substitution rate (K_s_) between *L. multiflorum* and closely related species.

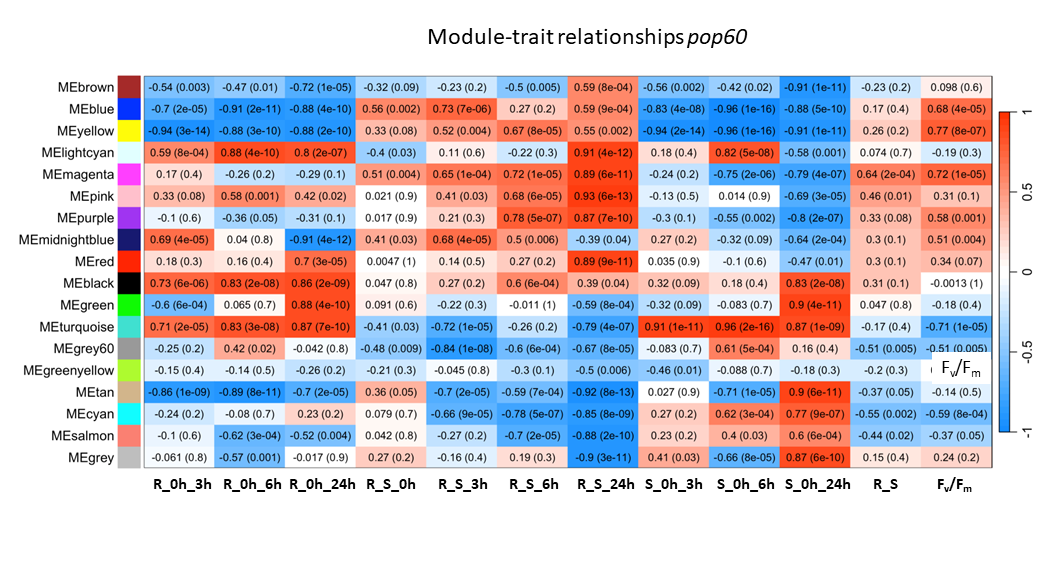

A

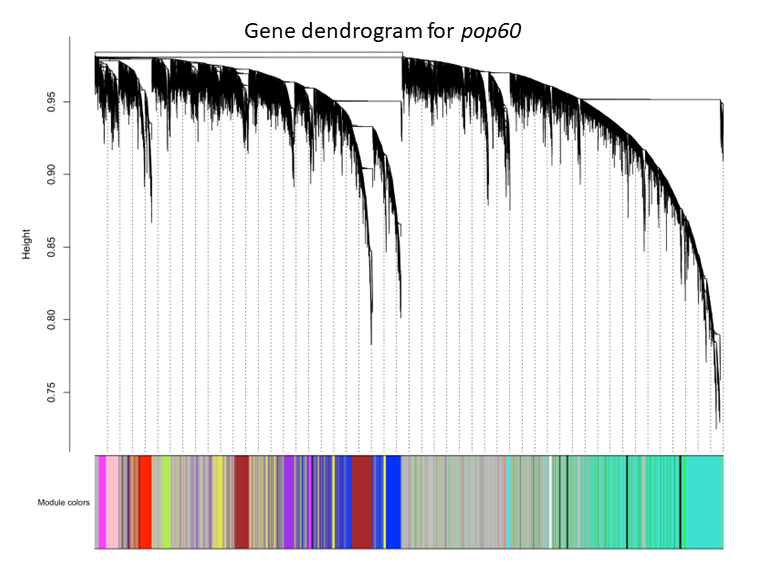

B

Fig. S2. Weighted gene co-expression network analysis of the F_3_ paraquat-resistant pop60 population. (A) Heat map of module-trait relationships shows a correlation from more negative (blue) to more positive (red) for each module, which are given names with different colors. Each column indicates a comparison between time points or individuals. (B) Hierarchical cluster trees show the co-expression modules identified by WGCNA.

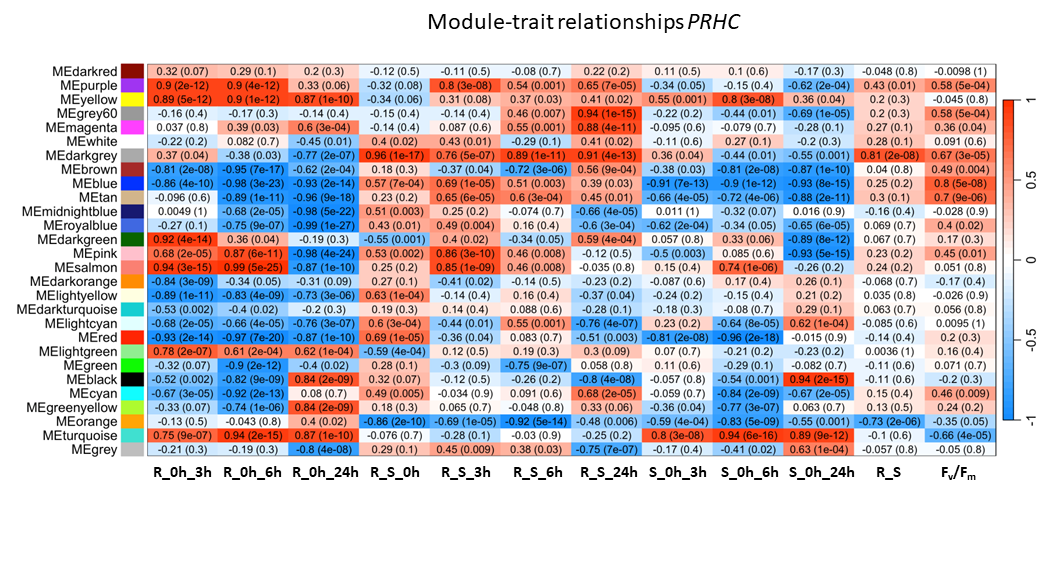

A

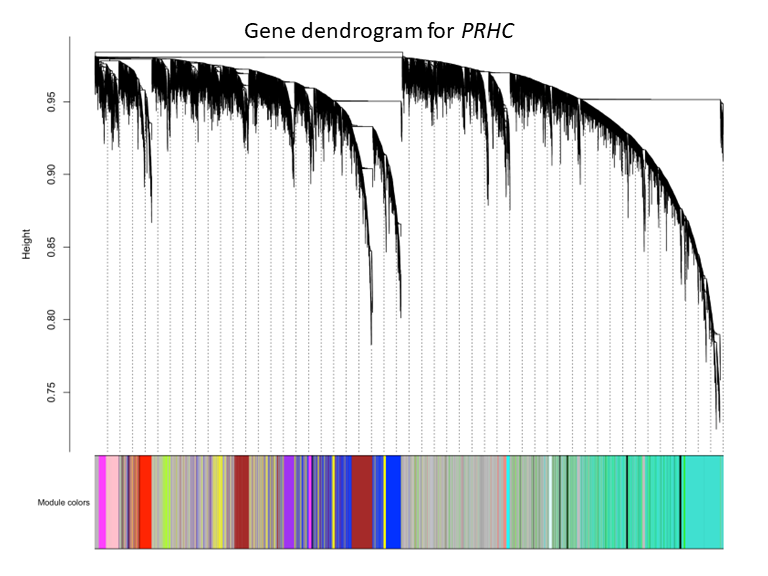

B

Fig. S3. Weighted gene co-expression network analysis of the F_3_ paraquat-resistant PRHC population. (A) Heat map of module-trait relationships shows relationship from more negative (blue) to more positive (red) of each module color. Each column indicates a comparison between time points or individuals. (B) Hierarchical cluster trees show the co-expression modules identified by WGCNA.

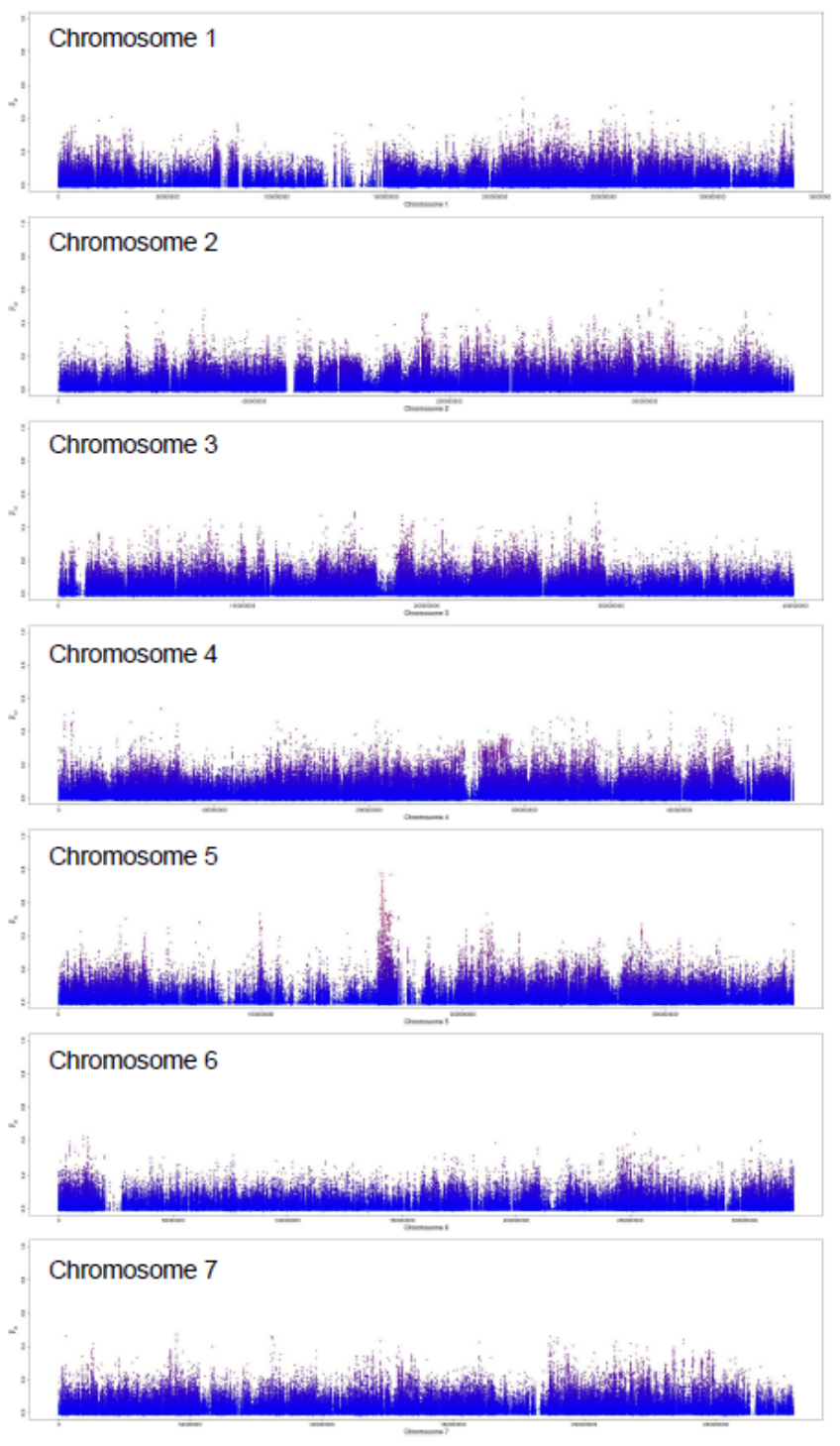

Fig. S4. Genome scan of Fst at all bi-allelic SNPs between paraquat-resistant and -susceptible populations of *L. multiflorum*.

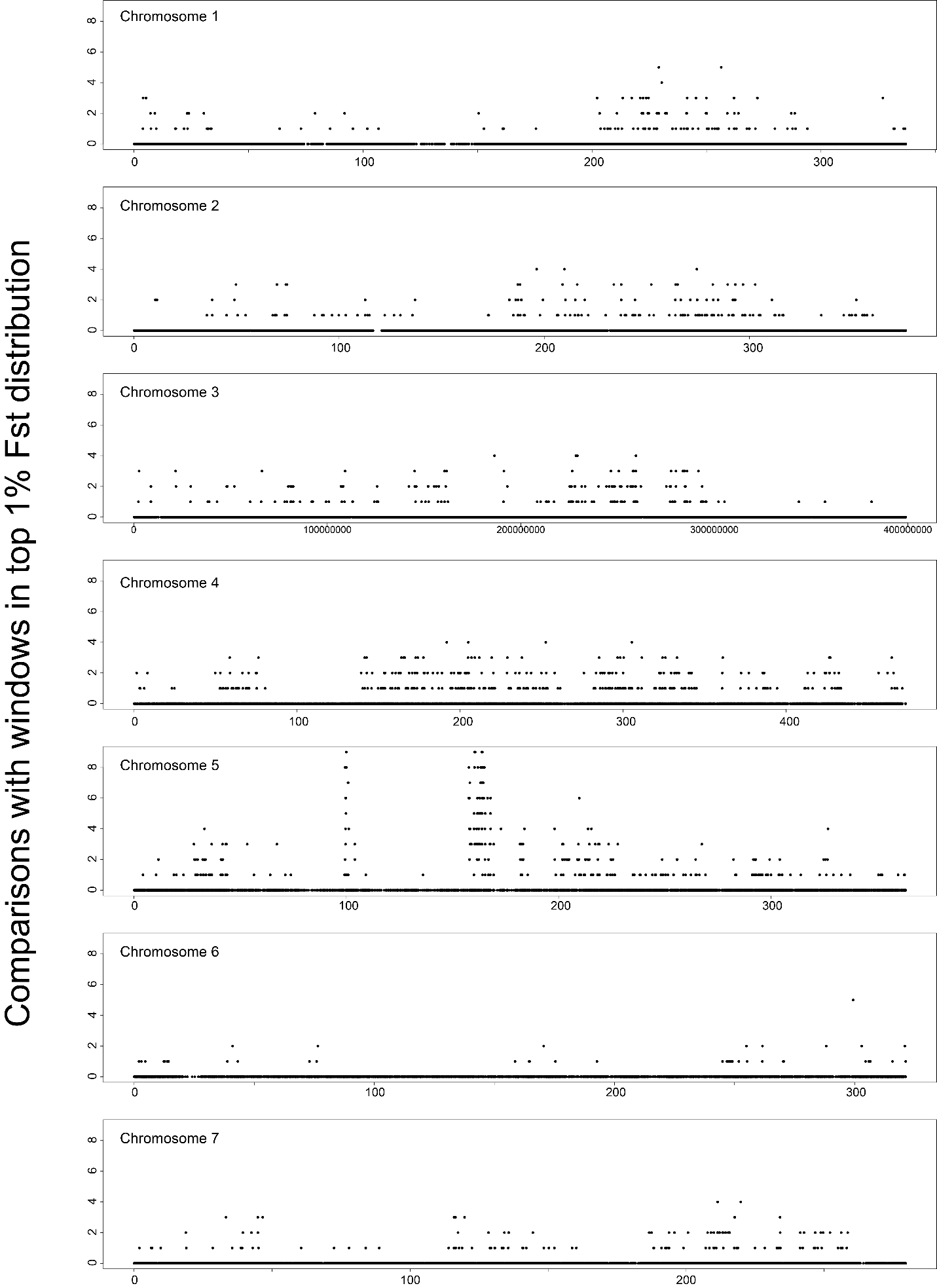

Fig. S5. The number of pairwise comparisons between resistant and susceptible populations where Fst (calculated in 25 kb windows) was found in the top 1% of the distribution across windows.

Table S1. Summary of *L. multiflorum* assembly and annotation.

| **Characteristics** | **Value** |
| --- | --- |
| Genome size (bp) | 2,855,700,136 |
| Number of contigs | 253 |
| N_50_ (bp) | 363,560,625 |
| L_50_ (scaffold) | 4 |
| L_90_ | 7 |
| Repetitive elements (%) | 82.61 |
| Average gene length (bp) | 4,752 |
| Number of annotated genes | 49,295 |
| BUSCO | Single-copy: 72.5%  Duplicated: 20.3%  Fragmented: 0.2%  Missing: 7.0% |

Table S2. Repetitive element content in the *L. multiflorum* genome.

| **TE Class** |  | **Count** | **%** |
| --- | --- | --- | --- |
| LTR |  |  |  |
|  | Copia | 149860 | 4.80 |
|  | Gypsy | 1135594 | 35.04 |
|  | Unknown | 1449067 | 23.72 |
| TIR |  |  |  |
|  | CACTA | 328256 | 4.67 |
|  | Mutator | 180065 | 2.58 |
|  | PIF_Harbinger | 112106 | 1.85 |
|  | Tc1_Mariner | 127098 | 1.20 |
|  | hAT | 40264 | 0.51 |
| nonLTR |  |  |  |
|  | LINE_element | 10310 | 0.19 |
|  | unknown | 216 | 0.01 |
| nonTIR |  |  |  |
|  | helitron | 195580 | 3.18 |
| repeat_region |  | 138747508 | 4.86 |
| Total |  | 2358935386 | 82.61 |

Table S3. Genomic regions identified in GWAS and number of annotated genes.

| **Pseudomolecule** | **Genomic coordinates** | **Annotated genes** |
| --- | --- | --- |
| 2 | 38,259,568-42,259,568 | 57 |
| 5 | 98,269,722-102,269,722 | 40 |
| 5 | 160,257,267-164,257,267 | 37 |
